# Optimized antibody with W32I mutation enhances antiviral efficacy against influenza

**DOI:** 10.1101/2025.01.24.634663

**Authors:** Shanshan Guan, Qingyu Wang, Jiaojiao Nie, Xin Yao, Kunpeng Xie, Weihao Zhao, Yaotian Chang, Lvzhou Zhu, Jiaru Hui, Tieyan Yin, Xiaopan Liu, Yaming Shan

## Abstract

The influenza virus has caused a global pandemic with significant morbidity and mortality, highlighting the need to optimize antibodies for improved antiviral efficacy. The 3E1 antibody effectively neutralizes influenza subtypes H1 and H5 by inhibiting acid-induced conformational changes of hemagglutinin (HA). This study aimed to optimize the antibody’s bioactivity by modifying amino acid residues, resulting in single-point mutants (3E1-L [W32I], 3E1-H [F103I]) and a double mutant (3E1-H+L [F103I, W32I]). The binding affinity, neutralizing activity, and antiviral mechanisms of the mutants were evaluated. Notably, the 3E1-L mutant showed significantly enhanced antiviral activity against H1N1 and H3N2 compared to wild-type 3E1, inhibiting both viral entry and release. The prophylactic and therapeutic efficacy of the 3E1-L mutant was validated. Molecular dynamics simulations of the 3E1-L/HA complex showed that the W32I mutation reduces steric hindrance between tryptophan at position 32 and the complementarity-determining region (CDR) L1 loop of HA. In conclusion, the W32I substitution enhances the antiviral activity of wild-type 3E1, making the optimization of 3E1-L a promising strategy for developing more effective influenza therapies.

## Introduction

The influenza virus (IV), a member of the *Orthomyxoviridae* family, is a highly contagious pathogen responsible for significant global morbidity and mortality each year. Seasonal IV cause approximately 3 to 5 million severe illness and 290, 000 to 650, 000 deaths annually (Lagacé-Wiens *et al*, 2010; Ledford, 2008; Liu, 2014). The primary strategy for preventing IV infections relies on vaccination. However, the effectiveness of seasonal influenza vaccines depends on how well the vaccine strains match the circulating virus strains. As a result, the protective efficacy of vaccines can vary from year to year (Kim *et al*, 2022; Schultz-Cherry & Jones, 2010).

In addition to vaccination, antiviral drugs, including chemicals and monoclonal antibodies, provide alternative options for treating influenza, particularly during the early stages of an IV pandemic or to control the spread of infection(Beigel & Bray, 2008; Erlich, 2019; Hurt *et al*, 2006). Current antiviral drugs for influenza mainly consist of neuraminidase (NA) inhibitors, such as Oseltamivir, Zanamivir, and Peramivir. These drugs primarily act by preventing the release of progeny viruses from infected cells (Kolocouris *et al*, 1994; Wu *et al*, 2016). However, the widespread use of these antivirals has led to the emergence of drug-resistant strains, which retain their transmissibility (Blick *et al*, 1998; Sarker *et al*, 2022). Given the lack of cross-protective vaccines and the growing issue of antiviral resistance, there is an urgent need to develop monoclonal antibodies with broad protective activity against influenza for both prevention and treatment (Loregian *et al*, 2014).

Several monoclonal antibodies targeting the head and stem regions of IV hemagglutinin (HA) have been discovered, with those targeting the stem region of HA showing broad-spectrum potential. Among these, antibodies such as C179 (Okuno *et al*, 1993), MEDI8852 (Kallewaard *et al*, 2016), and 3E1 (Wang *et al*, 2016) have demonstrated promising neutralizing activity. The discovery of the murine monoclonal antibody C179, which binds to the conserved region of the HA stalk, has shown that antibodies targeting this region can neutralize a wide range of influenza strains, including subtypes H1 and H2 (Okuno *et al*., 1993). The therapeutic potential of these broadly neutralizing antibodies is being actively explored, with the possibility of their mechanisms influencing the development of cross-protective vaccines for influenza (Corti & Lanzavecchia, 2013; Yewdell, 2013).

Wang et al. reported a human monoclonal antibody, 3E1, which binds to a conserved region slightly below the stalk of HA (primarily consisting of a short helix in HA2 and a portion of the fusion peptide). Both *in vivo* and *in vitro* studies have demonstrated that antibody 3E1 effectively neutralizes infections by IV subtypes H1 and H5 by inhibiting the acid-induced conformational change of the HA protein. Notably, 3E1 has been shown to provide protection to mice exposed to H1N1 and H5N6 viruses. Although 3E1 exhibits broad neutralizing activity against multiple strains of the H1 and H5 subtypes, further optimization is needed to enhance its antiviral potency.

In this study, based on the structural analysis of the 3E1 antibody, the researchers selected two antibody residues near the HA binding interface—W32 on the light chain and F103 on the heavy chain—as potential mutation sites. Three mutants were designed: single-point mutations 3E1-L (W32I) and 3E1-H (F103I), and the double mutant 3E1-H+L (F103I, W32I). This approach aimed to optimize the antibody’s interaction with HA, potentially improving its antiviral activity.

The binding affinity of the mutants to HA and their inhibitory effects on the virus were assessed through enzyme-linked immunosorbent assays (ELISA) and microneutralization assays *in vitro*. Additionally, molecular dynamics simulations were used to investigate the microscopic interaction mechanisms between the mutant antibodies and HA. The findings from this research provide valuable insights and a theoretical foundation for the development of more effective, broadly neutralizing antibody therapies against influenza A viruses.

## Results

### 3E1-L increases reactivity against HA

To develop a highly effective HA-specific antibody, three mutants were designed: single-point mutants 3E1-L (W32I), 3E1-H (F103I), and a double mutant 3E1-H+L (F103I, W32I), all of which were expressed in ExpiCHO cells (**Fig. 1A**). The thermal stability of these antibodies was evaluated by measuring their melting temperatures under various solution conditions, as shown in **Fig. 1B**. The first transition temperature (Tm1) represents the lowest melting temperature, corresponding to the CH2 domain, while the second transition temperature (Tm2) represents the highest peak, corresponding to the Fab domain. The Tm1 and Tm2 values demonstrate that all mutants retain thermal stability.

**Figure 1.**
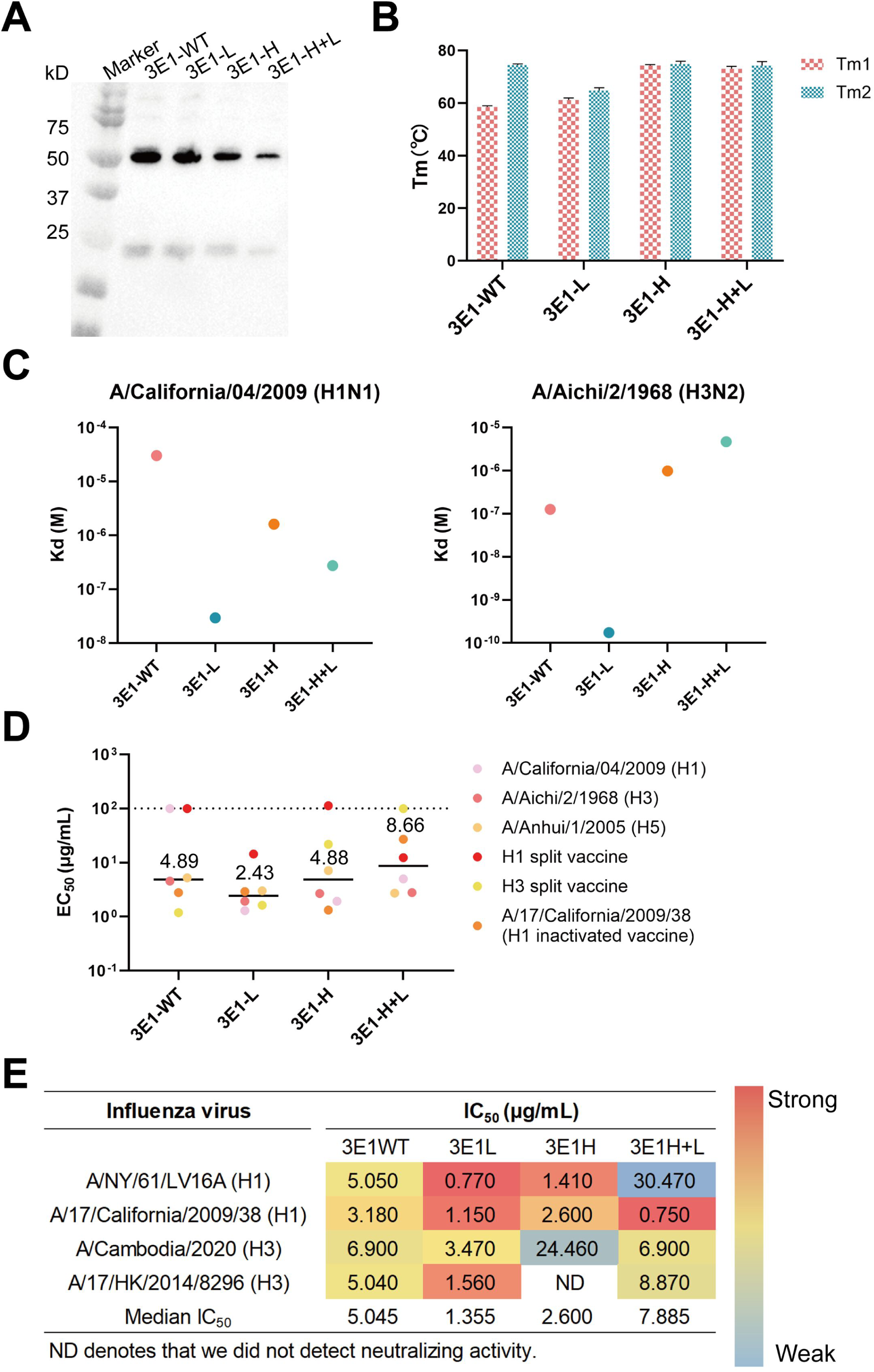
*In vitro* binding and neutralization activities of 3E1-WT and mutants. (A) SDS-PAGE analysis of the purified antibodies. (B) Histogram showing the Tm1 and Tm2 values for each antibody. (C) Binding affinity of the purified antibodies to HA proteins. (D) EC_50_ values of the purified antibodies binding to HA proteins and split virions, determined by ELISA. (E) IC_50_ values of the purified antibodies against H1 or H3 viruses, determined by microneutralization assay.

Next, the binding activities of the mutants to HA were assessed using bio-layer interferometry (BLI) and ELISA (**Fig. 1C, D**). BLI results revealed that 3E1-L exhibited enhanced affinity for HA compared to 3E1-WT. Specifically, 3E1-L showed a 2-fold increase in binding to both H1 and H3 subtypes of HA. To determine whether this improved binding translated into stronger antiviral activity, we measured the neutralizing potency of the antibodies in Madin-Darby Canine Kidney (MDCK) cells infected with H1N1 and H3N2 viruses. The 3E1-L antibody demonstrated potent neutralization, with a median IC_50_ of 1.355 μg/mL (range: 0.770∼3.470 μg/mL), compared to 3E1-WT, which had a median IC_50_ of 5.045 μg/mL (range: 3.180∼6.900 μg/mL) (**Fig. 1E**). Additionally, we observed that the co-mutations (3E1-H+L) did not yield a significant increase in potency over the single mutants, likely due to overlapping or redundant binding epitopes. These results indicate that optimizing the 3E1-L antibody led to a 4-fold increase in neutralizing potency and a 2-fold increase in binding affinity for the H1 and H3 subtypes of HA.

### 3E1-L generates antiviral activity through a dual mechanism

Several studies have revealed that HA stem antibodies reduce viral egress by blocking viral invasion, preventing low pH-induced membrane fusion, or inhibiting the cleavage of HA0 into HA1 and HA2 (Kosik *et al*, 2019; Sun *et al*, 2024). To confirm the mechanisms involved for the mutants, we first detected influenza A virus nucleoprotein (NP) antigen in the supernatants and lysates of MDCK cells infected with either H1N1 or H3N2 viruses to assess the ability to block viral invasion (**Fig. 2A, B**). Virus release inhibition analysis showed that viruses were still detected in the supernatants of all groups, except for the 3E1-L group, indicating that 3E1-L significantly inhibits the release of H1 and H3 viruses (**Fig. 2C**). Consistent with the reactivity against H1 and H3 HA, the dual mechanism of 3E1-L was observed against both H1N1 and H3N2 viruses, demonstrating that 3E1-L effectively blocks both viral invasion and release compared to 3E1-WT.

**Figure 2.**
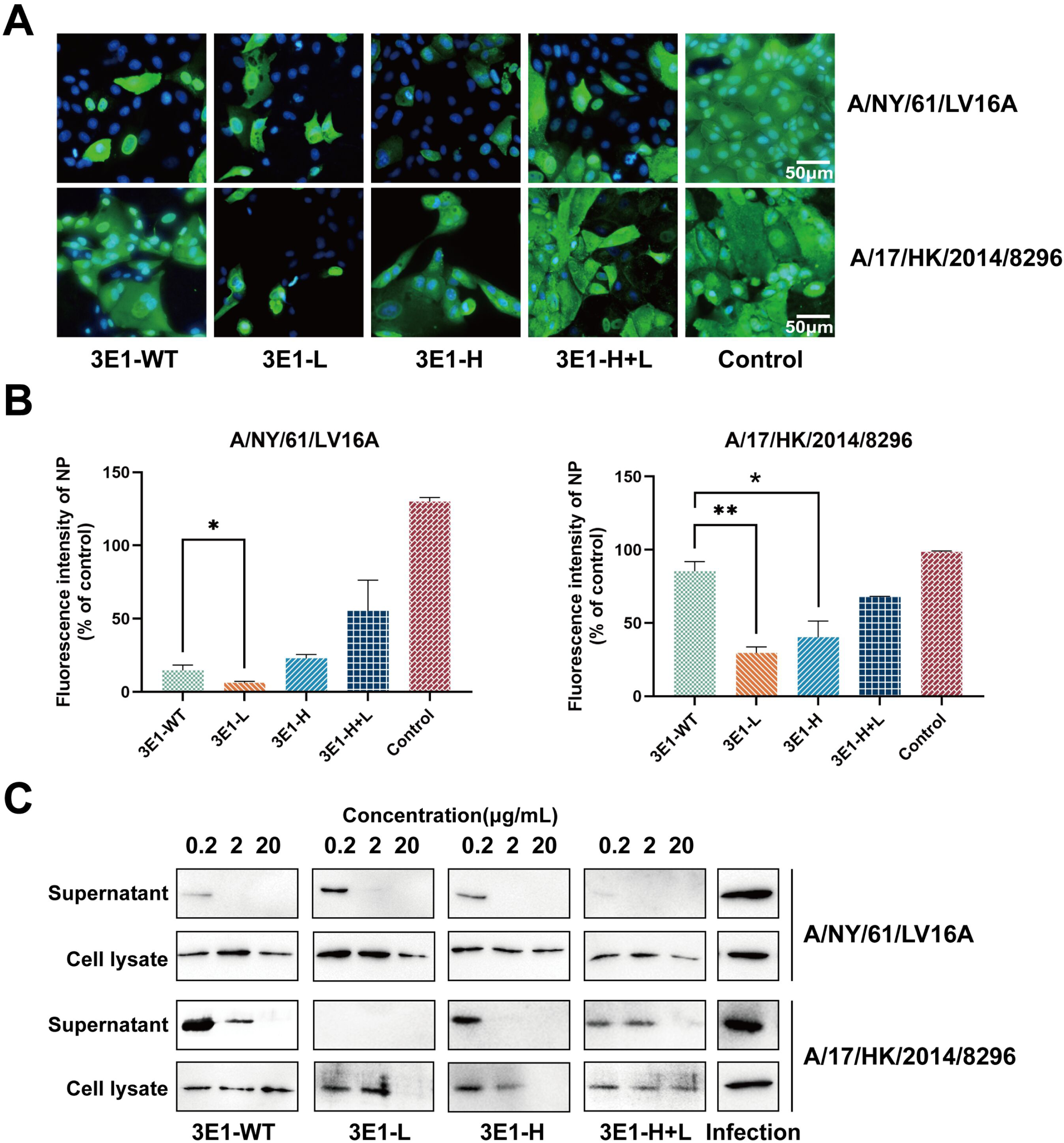
Inhibitory mechanisms of 3E1-WT and mutants. (A) Expression of influenza NP in MDCK cells following incubation with viruses preincubated with 3E1-WT or mutants, targeting A/NY/61/LV16A or A/17/HK/2014/8296, as indicated. (B) Quantitative analysis of NP fluorescence intensity. (C) Detection of influenza NP by western blotting in the supernatants and lysates of infected MDCK cells incubated with 3E1-WT or mutants.

Further mechanistic studies on the inhibition of HA0 cleavage and membrane fusion are presented in **Fig. S1**. The results indicate that 3E1-L maintains its capacity to inhibit HA-mediated membrane fusion and HA0 cleavage, akin to 3E1-WT. These findings further support the dual mechanism of action of 3E1-L.

### Prophylactic and therapeutic efficacy of 3E1-L in mice

The prophylactic and therapeutic efficacy of 3E1-L, based on its dual mechanism, was further evaluated. For prophylactic efficacy, mice were treated with either 3E1-WT or 3E1-L (1 mg/kg or 3 mg/kg) following a challenge with a lethal dose of A/NY/61/LV16A (H1N1) or A/17/HK/2014/8296 (H3N2) virus (**Fig. 3A**). Both antibodies provided full protection at the highest dose. At a dose of 1 mg/kg, 3E1-L (with 50%∼80% survival) offered more robust protection than 3E1-WT (with 0%∼50% survival), resulting in less weight loss (**Fig. 3B, C**). Additionally, lung viral titers in the 3E1-L group at 3 mg/kg were significantly lower than those in the 3E1-WT group at the same dose (**Fig. 3D**). Furthermore, lung tissue from 3E1-L-treated mice appeared nearly normal, with reduced leukocyte infiltration (**Fig. 3E**).

**Figure 3.**
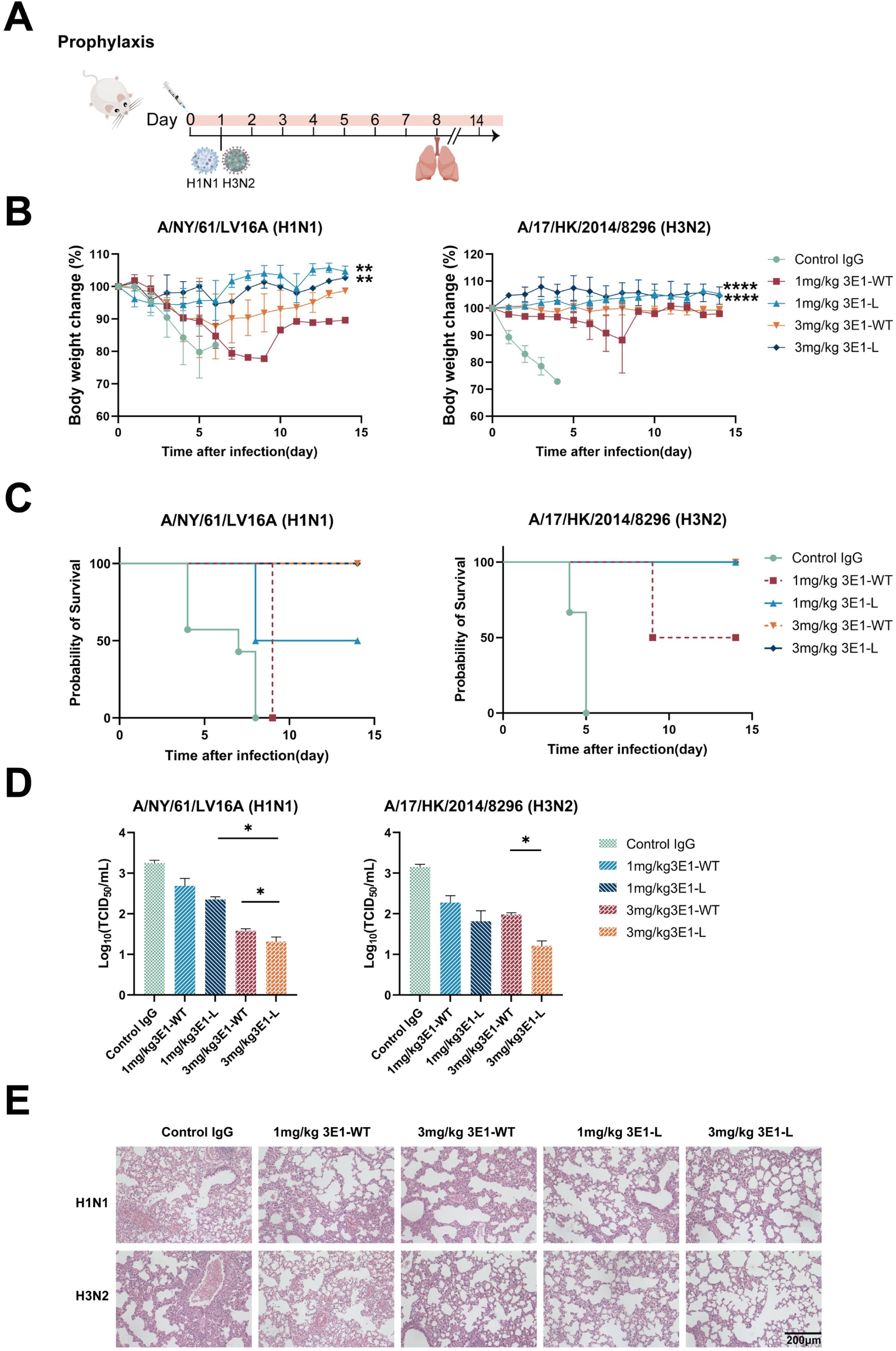
Prophylactic efficacy of 3E1-WT and 3E1-L in mice. (A) Experimental design for prophylactic testing. (B) Body weight changes (%) of mice (n=4/group) treated with 1, 3 mg/kg 3E1-WT or 3E1-L before lethal challenge with H1N1 and H3N2 viruses. (C) Survival curves of mice treated with 3E1-WT or 3E1-L at doses of 1 or 3 mg/kg before lethal challenge with H1N1 or H3N2 viruses. (D) Lung viral titers, determined by TCID_50_ assay. (E) H&E staining analysis of lung tissue from treated mice.

Based on the treatment efficacy of 3E1-L in mice, groups of mice were infected with a lethal dose of H1N1 or H3N2 virus and intravenously administered 3E1-WT or 3E1-L at a dose of 15 mg/kg (**Fig. 4A**). As shown in **Fig. 4B, C**, mice in the 3E1-L group exhibited slight weight loss, with 100% survival. In contrast, 3E1-WT only partially protected the mice, resulting in a survival rate of 30%. Additionally, lung viral titers in the 3E1-L group were nearly similar with 3E1-WT group (**Fig. 4D**). Consistent with these results, hematoxylin and eosin (H&E) staining analysis of the 3E1-L group revealed milder lesions (**Fig. 4E**).

**Figure 4.**
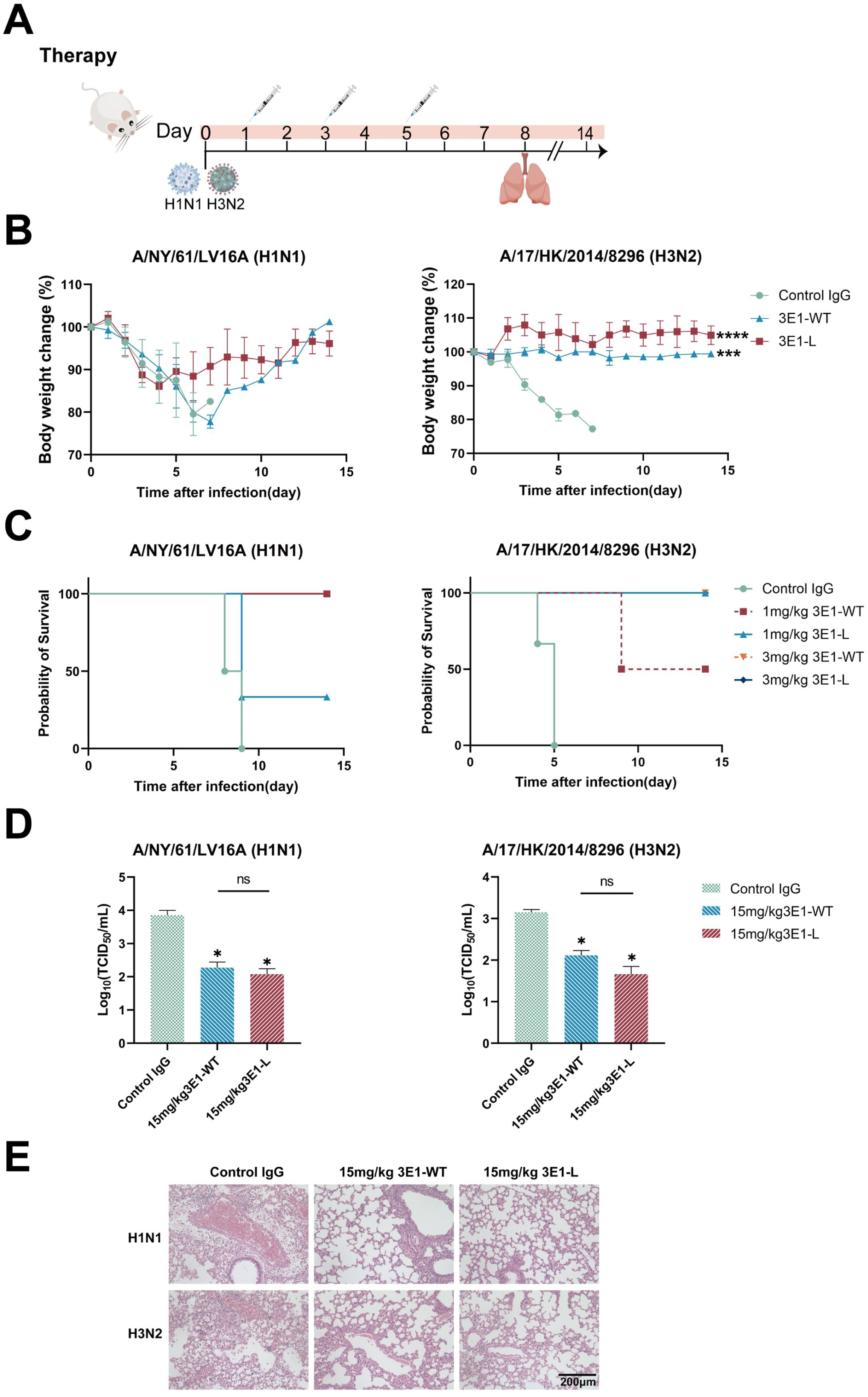
Therapeutic efficacy of 3E1-WT and 3E1-L in mice. (A) Experimental design for therapeutic testing. (B) Body weight changes (%) of mice (n=4/group) treated with 15 mg/kg 3E1-WT or 3E1-L following lethal challenge with H1N1 or H3N2 viruses. (C) Survival curves of mice treated with 15 mg/kg 3E1-WT or 3E1-L after lethal challenge with H1N1 and H3N2 viruses. (D) Lung viral titers, determined by TCID_50_ assay. (E) H&E staining analysis of lung tissue from treated mice.

These prophylactic and therapeutic results demonstrate that 3E1-L improves prophylactic efficacy and maintains treatment efficacy compared to 3E1-WT.

### Structure-based analysis of 3E1-L-mediated inhibition of H3 subtype HA

To elucidate the structure-based mediated inhibition of 3E1-L at the microscopic level, 100 ns molecular dynamics simulations and binding free energy calculations were conducted for both 3E1-WT and 3E1-L complex systems. During the entire simulation process, the changes in distance between each complementarity determining region (CDR) of the antibody and HA were monitored. As illustrated in **Fig.5**, there are notable differences in the examined distances between the wild-type and mutant systems. The W32I mutation in the antibody light chain results in reduced steric hindrance between the 32nd residue and HA, leading to a smaller and more stable distance between the CDRL1 loop and HA in the 3E1-L mutant compared to the wild-type system. Meanwhile, the distances between CDRL3, CDRH1, and CDRH2 and HA in the mutant system are also more stable compared to those in the 3E1-WT system.

**Figure 5.**
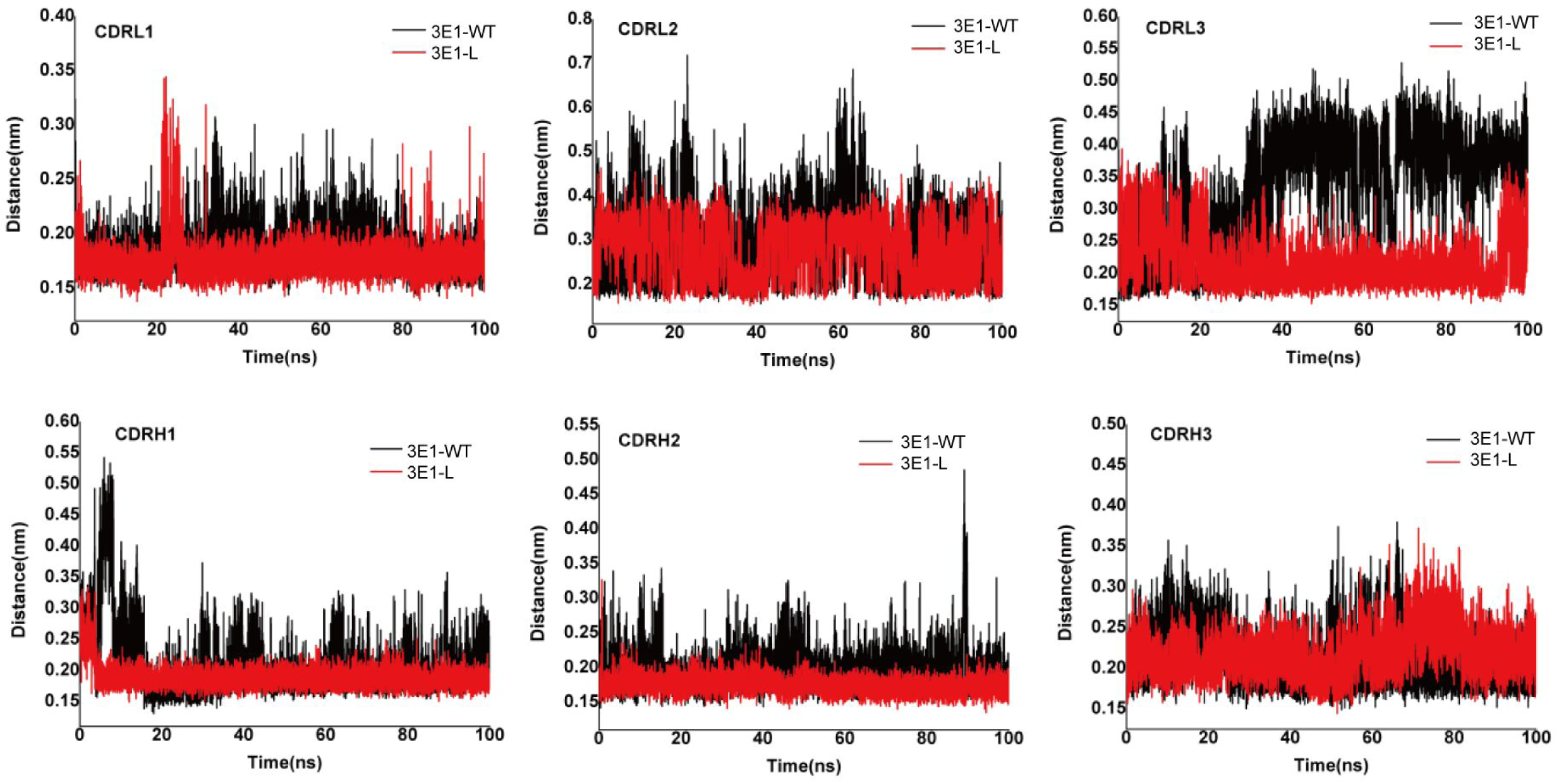
The distances between each CDRs of the antibody and HA in both 3E1-WT and 3E1-L complex systems.

The three-dimensional binding patterns reveal significant differences in the binding poses of both the 3E1-WT and mutants with HA (**Fig. 6**). In the 3E1-WT system, the antibody binding site is closer to the head region of HA, and the three heavy chain CDRs do not make sufficient contact with HA. In the 3E1-L system, the antibody binding site shifts slightly downward, and the W32I mutation reduces the steric hindrance between CDRL1 and HA, resulting in an increased effective contact area between CDRH1, CDRH2, and CDRH3 with the short helix of HA.

**Figure 6.**
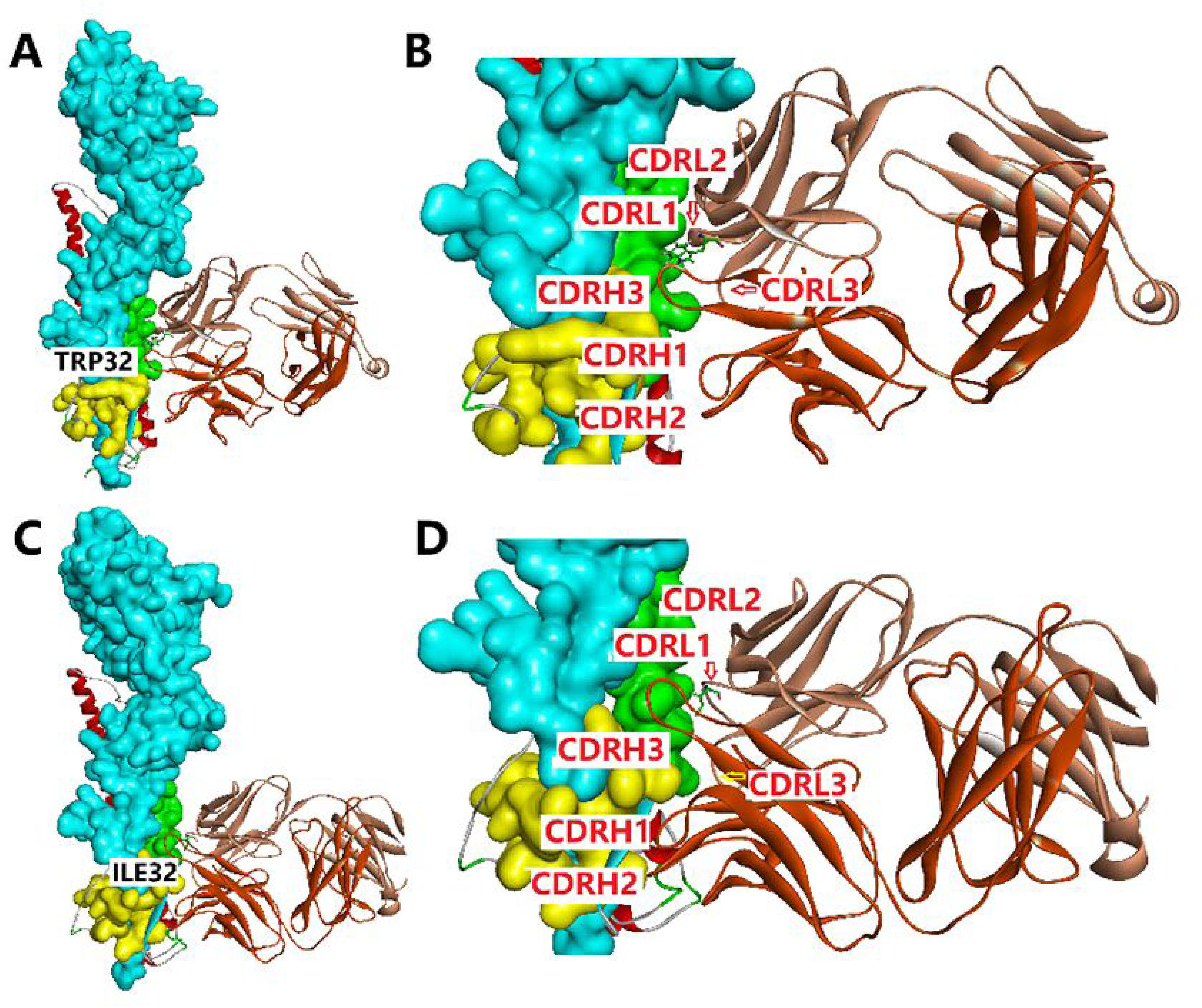
(A) The binding pose of 3E1-WT to the HA stem region. (B) The binding pose of CDRs of 3E1-WT to HA. (C) The binding pose of mutants to the HA stem region. (D)The binding pose of CDRs of mutants to the HA.

Based on the binding free energy results, as shown in **Fig. 7**, within the 3E1-L system, the binding energies between CDRL1, CDRL3, CDRH1, CDRH2, CDRH3, and HA are enhanced. Notably, CDRL3, CDRH1, and CDRH2 exhibit prominent binding contributions, suggesting that the mutations have altered the microscopic interactions between these regions and HA. Hydrogen bonding is a well-recognized primary force in antigen-antibody binding. The study examined the changes in the number of hydrogen bonds between the two systems and found that the number of hydrogen bonds in the 3E1-L system was significantly higher than that in the 3E1-WT system. Monitoring the number of hydrogen bonds between each CDR and HA reveals that the increase in hydrogen bonds in the mutated system primarily originates from the interactions between CDRL1, CDRH1, and CDRH2 with HA, as shown in **Fig. 8**.

**Figure 7.**
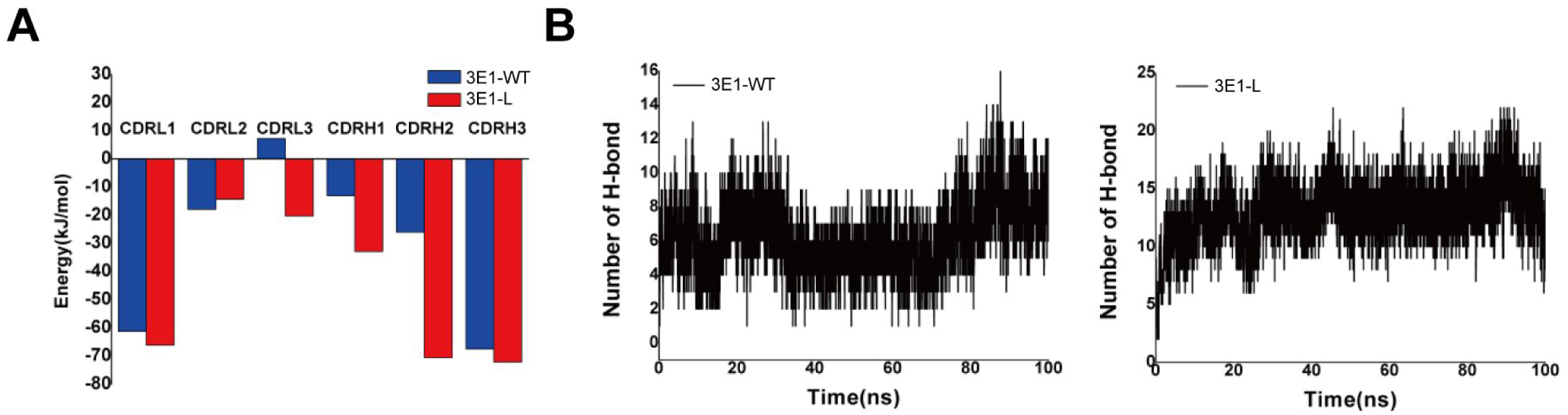
(A) The binding energy contribution of the CDRs in both 3E1-WT and 3E1-L complex systems. (B) The variation in the number of hydrogen bonds in both 3E1-WT and 3E1-L complex systems.

**Figure 8.**
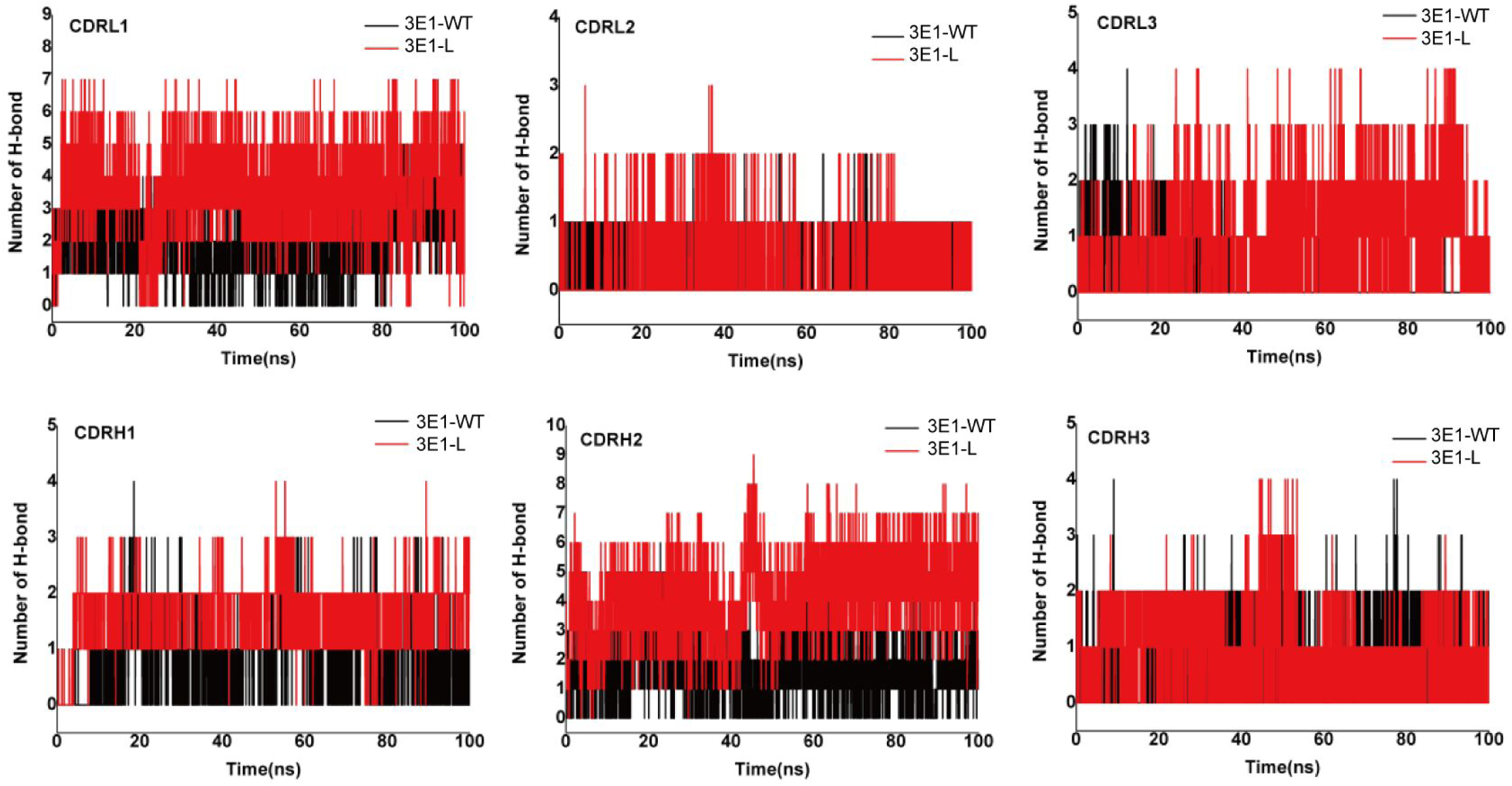
The variation in the number of hydrogen bonds between CDRs and HA in both 3E1-WT and 3E1-L complex systems.

## Discussion

IV presents a significant global health burden, impacting both public health and the socio-economic landscape. Currently, several promising strategies are under development, including neutralizing universal vaccines, small molecule drugs, and monoclonal antibodies. While various therapeutics have shown efficacy against respiratory viral infections, the emergence of highly variable strains poses a major challenge, making the development of broad-spectrum antiviral solutions essential.

Not all vaccinated individuals generate high-quality antibodies, such as the 3E1 antibody used in this study (Wang *et al*., 2016). However, engineering site-specific mutations in antibodies to enhance their antiviral activity, as exemplified by 3E1-L, may provide immediate and effective protection for a broader population. In this study, we demonstrate that a single amino acid substitution (Trp to Ile at position 32, W32I) in 3E1-L improves the antiviral efficacy of the original 3E1-WT antibody. Remarkably, 3E1-L exhibits a dual mechanism of action, inhibiting both viral entry and release, which contributes to its enhanced antiviral activity.

While the viral strains tested are limited, this strategy holds promise for improving the broad-spectrum efficacy of antibodies. 3E1-L has also shown potential as an effective preventive antibody in animal models, demonstrating rapid viral clearance in vivo. Furthermore, therapeutic efficacy remains high even after treatment. However, the strategy has been validated only at two dose levels of the mutant antibody. In the lower dose group, the advantages of 3E1-L were less pronounced, suggesting that further investigation using additional doses is needed to fully assess its therapeutic potential.

Dynamic analysis reveals that the W32I mutation in the antibody light chain significantly affects the binding properties of 3E1-L to the H3 HA, due to altered microscopic interactions between the antibody and HA. This mutation reduces steric hindrance between the 32nd residue and HA, resulting in a shorter distance between the CDRL1 loop of 3E1-L and HA. As a result, the binding stability of CDRL3, CDRH1, and CDRH2 with HA is enhanced. Additionally, the antibody binding site shifts slightly downward, increasing the effective contact area between CDRH1, CDRH2, and CDRH3 with the short helix of HA. The binding energies between each CDR and HA in 3E1-L are significantly increased, with CDRL3, CDRH1, and CDRH2 contributing most notably. Simulations show a significantly higher number of hydrogen bonds in the mutant system, primarily originating from interactions between CDRL1, CDRH1, and CDRH2 with HA.

With its robust anti-viral efficacy and potential for viral neutralization, 3E1-L offers an important prophylactic option in addition to vaccine approaches to protect vulnerable populations. Strong evidence from thorough characterization supports its potential to treat and prevent influenza in humans.

## Methods

### Viruses, cells, HA proteins and animals

A/NY/61/LV16A, A/17/California/2009/38, A/17/HK/2014/8296, A/Cambodia/E0826360/2020, were kindly provided by Changchun BCHT Biotechnology Co., Ltd. Influenza vaccines were obtained from Changchun Institute of Biological Products. MDCK cells (ATCC, USA) were cultured in Dulbecco’s Modified Eagle Medium (DMEM) supplemented with 10% fetal bovine serum (FBS) (Gibco, USA). ExpiCHO cells (Thermo Fisher Scientific, USA) were cultured in ExpiCHO medium (Gibco, USA). The baculovirus-expressed HA proteins of H1N1 A/California/04/09, H3N2 A/Aichi/2/1968, and H5N1 A/Anhui/1/2005 were purchased from Sino Biological (China).

6- to 8-week-old female specific pathogen-free BALB/c mice (weighing 18∼20 g) were purchased from Liaoning Changsheng Biotechnology Co., Ltd (Liaoning, China). Animal trials in this study were carried out in accordance with the Regulations for the Administration of Affairs Concerning Experimental Animals approved by the State Council of People’s Republic of China. All animal procedures were approved by the Institutional Animal Care and Use Committee of Jilin University (Permit number: YNPZSY2023096).

### Expression and purification of antibodies

The antibody sequence was synthesized by GenScript Biotech Co., Ltd. (NanJing, China) and inserted into the pVAX1 plasmid. Following transfection of the expression plasmids into ExpiCHO cells, antibody expression was carried out over a 6-day period. The antibody was then purified from the cell culture supernatants using Protein A affinity chromatography (Huiyan Biotechnology Co., Ltd., China).

### Western blotting

Sodium dodecyl sulfate-polyacrylamide gel electrophoresis (SDS-PAGE) using a 12% polyacrylamide gel was performed to analyze the antibodies. The gel was subsequently stained with Coomassie Blue (Sigma-Aldrich, St. Louis, MO, USA), followed by Western blotting using goat anti-human IgG conjugated to horseradish peroxidase (HRP; Solarbio, China) to identify the purified antibodies. The membrane was developed using the BeyoECL Moon kit (Beyotime Biotechnology, China) and visualized with the Tanon-5200 Chemiluminescent Imaging System (Tanon Science and Technology, China).

### ELISA

0.2 μg/well of HA proteins (Sino Biological, China) or split virions vaccines were coated into 96-well plates and incubated at 4℃ overnight, respectively. The 96-well plates were washed with phosphate buffered saline with Tween 20 (PBST) and blocked with 5% bovine serum albumin (BSA) at 37℃ for 2 h. Then, the plates were washed twice with PBST. Serial dilutions of serum with PBS were added to the 96-well plate and incubated at 37℃ for 1 h. After being washed four times with PBST, HRP-conjugated goat anti-mouse IgG (Dingguo Inc., China) with 1% BSA in PBS was added to the 96-well plate and incubated at 37°C for 45 min. After four washes with PBST, 3, 3, 5, 5-tetramethylbenzidine (TMB; TRANSGEN Biotech, China) color development solution was added to the 96-well plate at 100 μL/well and incubated for 20 min at room temperature in the dark. The reaction was stopped with 2M H_2_SO_4_, the absorbance at 450 nm was determined by an iMarK™ microplate reader (Bio-Rad, USA).

### *K*_d_ determination

To assess the binding affinity of antibodies, equilibrium dissociation constant (*K*_d_) values were determined using BLI with an OCTET RED96e (FortéBio, Germany). Antibodies at a concentration of 20 μg/mL were loaded onto anti-human Fc capture sensors and incubated with serial dilutions of HA proteins. Real-time kinetic analysis was performed, following the steps: Baseline (150 s), Loading (600 s), Baseline 2 (300 s), Association (700 s), and Dissociation (900 s). The *K*_d_ value was calculated as the ratio of the off-rate (*K_off_*) to the on-rate (*K_on_*).

### Microneutralization assay

To assess the neutralizing activity of antibodies, a microneutralization assay was performed using MDCK cells and 100 TCID_50_ (50% tissue culture infectious doses) of IV, as previously described (Gross *et al*, 2017). Briefly, 100 TCID_50_ of the virus was mixed with serial two-folded dilutions of purified antibodies and incubated at 37℃ for 2 hours. The mixture was then added to MDCK cells and incubated at 37℃ with 5% CO_2_ for 18∼20 hours. After incubation, the medium was removed from the microtiter plates, and the wells were washed with PBS. Next, 80% acetone was added to each well and incubated at 37℃ for 15 minutes. Anti-influenza A NP monoclonal antibody and HRP-conjugated goat anti-human IgG were used as the primary and secondary antibodies, respectively, in the ELISA.

### Cell-based entry inhibition assay

To evaluate the entry inhibition activity against influenza, the assay was performed as previously described (Dreyfus *et al*, 2012). Briefly, viruses were pre-incubated with antibodies at the indicated concentrations before being added to MDCK cells plated in infection medium (DMEM supplemented with 2% BSA and 20 μg/mL trypsin). After the virus and antibody mixture was added, the medium was removed and replaced with maintenance medium (DMEM supplemented with 2% BSA). The cells were then cultured for 16∼18 hours at 37℃ with 5% CO_2_. Following incubation, the supernatant was discarded, and the cells were fixed with 4% paraformaldehyde (PFA) solution. The cells were then sequentially incubated with influenza A NP antibody (Sino Biological, China) and an Alexa488-conjugated anti-rabbit secondary antibody (Abcam, USA), followed by 4’,6-diamidino-2-phenylindole (DAPI) staining. Finally, the cells were analyzed using an inverted fluorescence microscope (Nikon Instruments, USA).

### Antibody inhibition of viral release

To assess antibody-mediated inhibition of viral release, the assay was performed as previously described (Shen *et al*, 2017). The plated MDCK cells were infected with viruses at a 2-multiplicity of infection (MOI) in infection medium for 4 hours. After infection, the cells were incubated with serial dilutions of antibodies for 18 hours at 37°C with 5% CO_2_. The cell supernatants were then removed, and the cells were washed three times with PBS to remove extracellular viruses. Following this, the cells were re-incubated with different concentrations of antibodies for an additional 18 hours. A respiratory syncytial virus (RSV)-unrelated antibody was used as a control IgG. Viral release was assessed by western blotting of both the supernatants and cell lysates to quantify the virus present.

### Prophylactic and therapeutic efficacy studies in mice

To evaluate the prophylactic and therapeutic efficacy against IV infection *in vivo*, the following experimental procedures were used. In the prophylactic studies, 6- to 8-week-old female mice were intravenously injected with antibodies at doses of 1 mg/kg or 3 mg/kg. An RSV-unrelated antibody was used as a control IgG. On the second day post-injection, the mice were anesthetized with 0.5% pentobarbital sodium and challenged intranasally with 25 EID_50_ of either A/NY/61/LV16A or A/17/HK/2014/8296. Mice weights were monitored daily for up to 14 days following the challenge. At 8 days post-infection, lungs were collected for virus titration and histopathological analysis via H&E staining.

In the therapeutic studies, 6- to 8-week-old female mice were infected with 25 EID_50_ of either A/NY/61/LV16A or A/17/HK/2014/8296. Mice were then treated with antibody at a dose of 15 mg/kg at 1, 3, or 5 day post-infection. Lungs were collected for virus titration and H&E staining at 8 days post-infection. The weight of the mice was continuously monitored for up to 14 days after infection.

### Molecular dynamics simulation of the antigen-antibody complexes

The antigen-antibody complex systems were subjected to molecular dynamics simulation in periodic boundary condition using the Gromacs 5.1.5 software package with simple point charge water model (Hess & van der Vegt, 2006; Tiwari *et al*, 2022; Van Der Spoel *et al*, 2005). The Gromos 54 A7 force field was applied to describe both the antigen and antibody (Huang *et al*, 2011). First, energies of the complex systems were relaxed with steepest-descent energy minimization to eliminate steric clashes or incorrect geometry. Thereafter, 100ps Constant number of particles, Volume, and Temperature (NVT) and Constant number of particles, Pressure, and Temperature (NPT) were alternately operated with position restraints on the antigen and antibody to make the relaxation of the solvent molecules in two phases. The solvent molecules were equilibrated with the fixed protein at 310K, and the initial velocities were chosen from a Maxwellian distribution. Subsequently, the antigen and antibody were relaxed in a stepwise fashion and heated to 310K. The long-range electrostatic interactions were described with the particle mesh Ewald algorithm, with an interpolation order of 4, a grid spacing of 0.16 nm, and the Coulomb cutoff distance of 1.0 nm. Temperature and pressure coupling types were set with V-rescale and Parrinello-Rahman, respectively (Qian *et al*, 2016).

In the NVT ensemble, the temperature of the systems reached a plateau at the desired value (reference temperature=310K; time constant=0.1 ps). In addition, the equilibration of pressure (reference pressure=1.0 bar; time constant=2.0 ps) was performed under the NPT ensemble. The equilibrated ensembles were subjected to molecular dynamics simulations conducted for 100 ns employing the LINear Constraint Solver (LINCS) and SETTLE algorithm for bond constraints and geometry of water molecules. The 100ns molecular dynamics simulations were initiated for collecting data with a time step of 2 fs and coordinates were saved every 2 ps (Guan *et al*, 2018).

## Statistical analysis

Statistical analysis was performed using GraphPad Prism 8.0 (GraphPad Software, USA). The results are expressed as the mean ± standard deviation (SD). Ordinary one-way ANOVA was used to compare differences between groups. Statistical significance was determined by Tukey’s multiple comparisons test unless otherwise stated (indicated as follows: *, *P* < 0.05; **, *P* < 0.01; ***, *P* < 0.001; ns, not statistically significant).

## Acknowledgments

This work was supported by the Jilin Province Science and Technology Development Projects (Grant number: YDZJ202301ZYTS341), and the Youth Program of the National Natural Science Foundation of China (Grant number: 31901062).

## Author contributions

Conceptualization: SSG, QYW, JJN; Methodology: SSG, QYW, YMS; Investigation: XY, KPX, WHZ, YTC, LZZ, JRH, XPL, TYY; Visualization: YMS; Supervision: SSG, YMS; Writing-original draft: SSG, QYW; Writing—review&editing: SSG, QYW, YMS

---

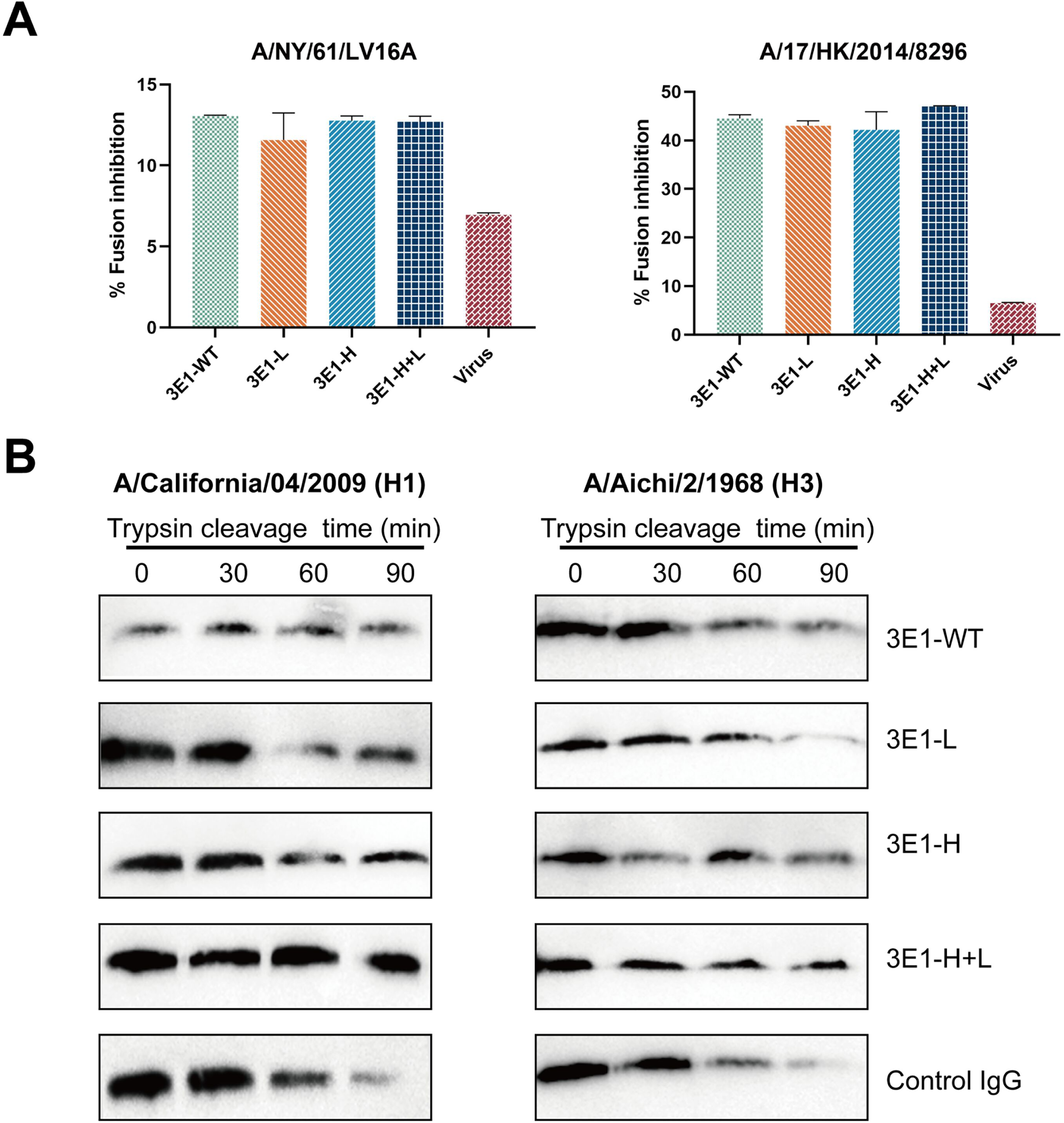

